# Acute Viral Infection Accelerates Neurodegeneration in a Mouse Model of ALS

**DOI:** 10.1101/2024.09.17.613525

**Authors:** Art Marzok, Jonathan P. Mapletoft, Braeden Cowbrough, Daniel B. Celeste, Michael R. D’Agostino, Jann C. Ang, Andrew T. Chen, Vithushan Surendran, Anna Dvorkin-Gheva, Ali Zhang, Hannah D. Stacey, Mannie Lam, Yasmine Kollar, Kevin R. Milnes, Sam Afkhami, Matthew S. Miller

**Author notes:** Denotes first author (Art Marzok, Jonathan P. Mapletoft). Denotes corresponding author.

## Abstract

While several viral infections have been associated with amyotrophic lateral sclerosis (ALS), the mechanism(s) through which they promote disease has remained almost entirely elusive. This study investigated the impact of common, acute viral infections prior to disease onset on ALS progression in the SOD1^G93A^ mouse model. A single sublethal infection prior to onset of ALS clinical signs was associated with markedly accelerated ALS disease progression characterized by rapid loss of hindlimb function. Prior infection resulted in gliosis in the lumbar spine and upregulation of transcriptional pathways involved in inflammatory responses, metabolic dysregulation, and muscular dysfunction. Therapeutic suppression of gliosis with an anti-inflammatory small molecule, or administration of a direct-acting antiviral, was associated with significantly improved ALS clinical signs, akin to what was observed in uninfected animals. This study provides causal and mechanistic evidence that the immune response elicited by acute viral infections may be an important etiological factor that alters ALS disease trajectory, and provides insight into novel therapeutic and preventative strategies for ALS.

**One Sentence Summary:** Acute viral infection with influenza A virus and SARS-CoV-2 accelerates the progression of ALS in SOD1^G93A^ mice.

## Introduction

Amyotrophic Lateral Sclerosis (ALS) is an incurable motor neuron disease which is characterized by progressive deterioration of upper and lower motor neurons leading to muscle wasting and paralysis^1,2^. The disease is remarkably heterogeneous in age-of-onset and in presentation of symptoms, even for individuals harboring the same ALS-associated mutations^3–6^. A combination of genetic and environmental factors are therefore believed to play critical roles in disease development and progression^6^.

Viruses have been characterized as environmental risk factors for ALS and other neurodegenerative diseases^7^. However, several of these associations have been erroneous and study designs have led to an enrichment bias favoring viruses that cause chronic/latent and neurotropic infections^8^. Poliovirus was the first virus associated with ALS due to its neurotropism and ability to cause post-poliomyelitis which phenotypically resembles ALS^7^. Subsequent studies have demonstrated that the enterovirus-associated 3C protease is able to cleave TAR DNA binding protein 43 (TDP-43), a pathophysiological feature observed in 97% of ALS cases^9,10^. Additionally, a sublethal intracerebroventricular infection with the coxsackie virus B3, a type of enterovirus, accelerated ALS disease progression in SOD1^G85R^ mice^11^. There has also been compelling evidence linking endogenous retroviruses such as Human Endogenous Retrovirus-K (HERV-K) to ALS, as transgenic mice constitutively expressing the *env* protein of HERV-K develop symptoms of progressive motor neuron disease^12,13^. Such observations have supported on-going clinical trials evaluating antiretroviral therapy in ALS patients^14–16^.

Although common, acute non-neurotropic viral infections may also be associated with ALS and other neurodegenerative diseases, they have been widely understudied since ubiquitous infections, especially those that occur significantly prior to the onset of disease (i.e. “hit-and-run”), are difficult to associate using conventional epidemiological approaches. This, is due, in part, to the fact that these viruses are likely cleared long before ALS symptom onset begins and seropositivity is high amongst the general population. Elegant recent work using data from biobank repositories across multiple countries showed an increased risk of developing neurodegenerative diseases such as ALS from acute viral infections, with increased hazard ratios persisting even 15 years following infection^17^. However, much remains to be understood regarding the mechanism through which viral infections impact ALS.

Influenza A viruses (IAV) are a group of single-stranded RNA viruses which predominantly result in acute, self-limiting respiratory infections^18^. Infection is associated with systemic elevation of pro-inflammatory soluble mediators which can potentiate neuroinflammation. Indeed, non-neurotropic IAV strains have been shown to directly induce long-lasting neuroinflammation with accompanying decreases in dendritic spine density to hippocampal neurons – a typical hallmark of neurodegeneration^19^. The COVID-19 pandemic, caused by SARS-CoV-2, has also led to lasting neurological manifestations during “long COVID”, with studies showing long-term neurological sequelae in approximately 33% of recovered COVID-19 patients, even 6 months post-infection^20,21^. In addition, SARS-CoV-2 RNA and antigens have been found in the CNS of COVID-19 patients^22^. Strikingly, SARS-CoV-2 infection has already been associated with increased susceptibility to Parkinsons disease as well as other neurodegenerative diseases^23,24^. In addition to inducing pathological immune responses, viral infections have been shown to perturb many of the same pathophysiological pathways underlying motor neuron death in ALS^8^.

A common mechanism through which diverse viral infections may exacerbate ALS is through the induction of aberrant host immune responses that can lead to neuroinflammation^25^. Inflammation is a hallmark characteristic of both viral infections and ALS, with elevated peripheral inflammation and neuroinflammation positively correlating with disease severity^26^. ALS patients often experience heightened levels of gliosis and pro-inflammatory cytokines in the peripheral blood and cerebral spinal fluid (CSF)^27^. Given this close interplay between inflammation and ALS, common acute viral infections may exacerbate ALS disease progression through further stimulating the immune system and elevating pre-existing CNS and systemic inflammatory mediators. However, many studies suggesting an association between ALS and prior infection have been correlative and lack mechanistic insights as to how viral infections potentiate disease. Furthermore, it remains unclear whether common acute respiratory infections impact ALS disease progression, and the mechanism(s) through which this might occur.

Here, we demonstrate that a single acute infection with either IAV or SARS-CoV-2 during the pre-clinical stages of ALS disease significantly accelerates ALS progression in SOD1^G93A^ mice. Interestingly, intramuscular administration of inactivated IAV resulted in similar disease acceleration, a phenomenon not observed following intranasal delivery. Mechanistically, this acceleration of ALS progression was accompanied by activation of microglia and astrocytes in the lumbar spinal cord of SOD1^G93A^ mice following infection, and transcriptomic enrichment of pathways relating to inflammation, immunity, and muscular dysfunction. Importantly, our findings further reveal that the inhibition of microglial activation during infection, or treatment with antivirals, helps mitigate ALS acceleration in these mice. This study thus provides direct causal and mechanistic evidence of acute respiratory infections exacerbating ALS progression in a mouse model of disease.

## Results

### Acute IAV infection prior to ALS onset accelerates disease progression in SOD1^G93A^ mice

Despite over 30 years of reported associations between viral infections and ALS, no prior studies have systematically explored the role of acute, non-neurotropic viral infections on the progression of ALS. To address this critical gap, we used influenza A virus (IAV) as a model given its ubiquity. Indeed, most individuals have experienced their first influenza infection by five years of age, and are re-infected regularly throughout the course of their life^28–30^. To determine whether IAV infection during the pre-clinical stages of ALS disease accelerates the onset and/or progression of ALS, wildtype BL6SJL (WT) and B6SJL-TgN(SOD1*G93A)1Gur (SOD1^G93A^) mice were infected intranasally (i.n.) with a sublethal dose (300 PFU) of mouse-adapted influenza A virus (strain A/Puerto Rico/8/1934 H1N1, (PR8)) at day 60 of age (**Fig. 1A**). This age was chosen based on the well-characterized kinetics of disease onset in this model as reported in the literature (typically, early onset is detectable at day 90)^31^. Indeed, there were no statistical differences observed between WT and SOD1^G93A^ mice in rotarod performance at day 60 of age (mean 51.1 and 47.9 seconds), indicating no observable phenotypic signs of ALS onset in SOD1^G93A^ mice (**Fig. 1B**). Following infection, no infectious virus was detected in lung and brain homogenates of WT or SOD1^G93A^ mice at 14 days post-infection (dpi) (**Supplemental Fig. 1A and 1B**). This is consistent with previous literature demonstrating that infection with this strain of IAV is acute and non-neurotropic^32^. After recovering from infection, mice were tested on the rotarod apparatus to measure motor function and general coordination and were compared to uninfected controls. Mice were tested weekly and the latency for falling off the rotating rod was recorded. Previously infected SOD1^G93A^ mice performed significantly worse on the rotarod test, as measured by the weeks elapsed to reach 50% of their initial fall time, when compared to uninfected SOD1^G93A^ mice (Mean 9.137 weeks and 11.182 weeks, p = 0.0002) (**Fig. 1C**). In addition, these mice reached endpoint due to ALS clinical signs, as defined by complete hind-limb paralysis and loss of the righting reflex, significantly faster than uninfected SOD1^G93A^ mice (**Fig. 1D**). Taken together, these data demonstrate that a single, acute, sublethal IAV infection during the pre-clinical stages of ALS significantly accelerates disease progression.

**Figure 1.**
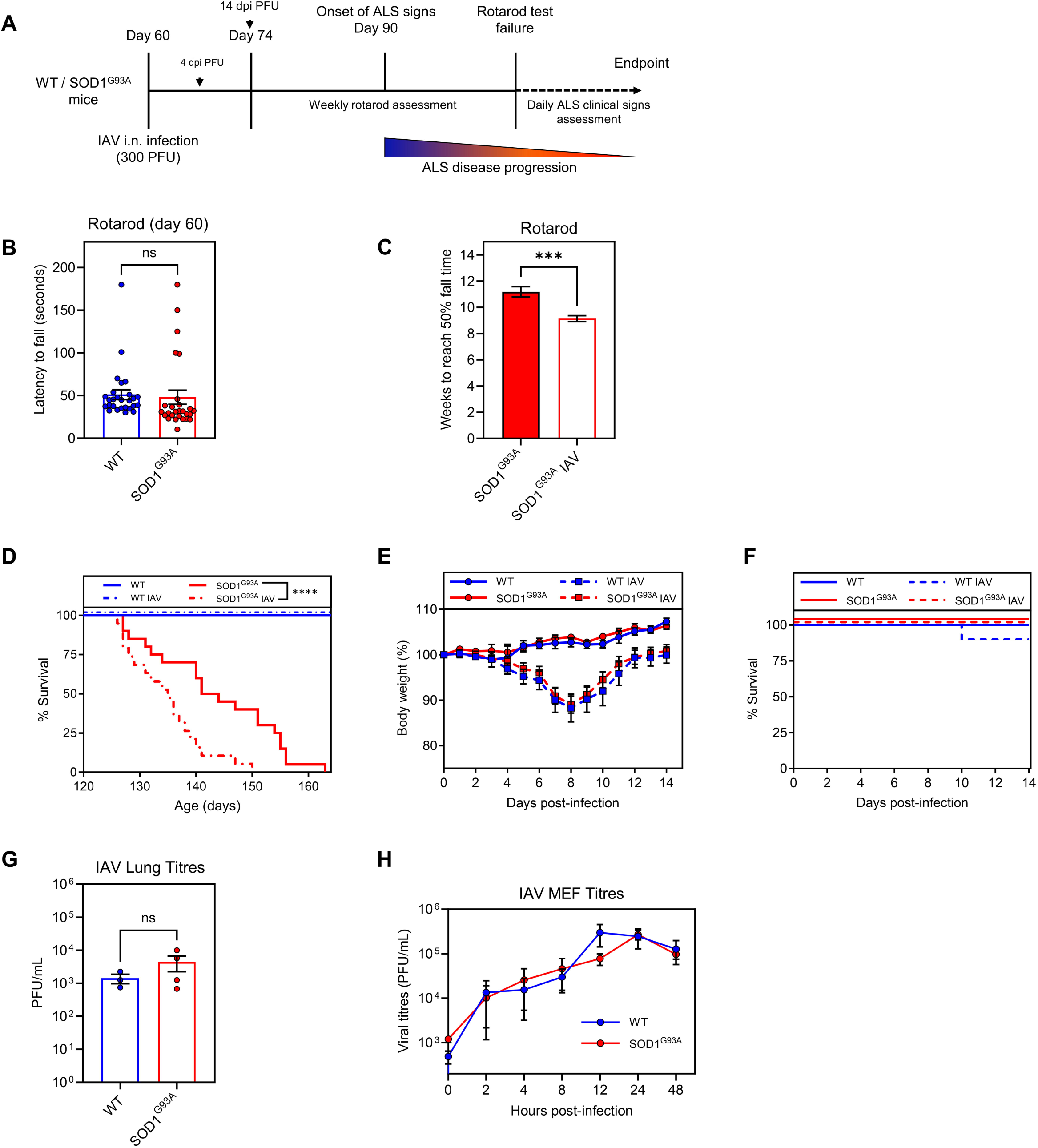
Acute IAV infection prior to ALS onset accelerates disease progression in S0D1G93Amice. **A)** Schematic of the experimental timeline for IAV infection of mice. **B)** Raw rotarod test fall times of wildtype (WT) (n=9) and SOD1893A(n=9) mice at day 60 of age. **C)** Weeks for IAV infected (n=14) and control (n=11) SOD1893A mice to reach 50% of their initial fall time. **D)** ALS-related endpoint (survival} of uninfected WT (n=9) and SOD1893A (n=20) and IAV infected WT (n=8) and SOD1893A(n=19) mice. **E)** Weight loss following IAV infection of WT (n=10) and SOD1893A(n=19) mice as compared to mock-infected WT (n=8) and SOD1893A(n=10) mice. **F)** Survival of mice following IAV infection. **G)** Lung viral tilers 4 days post-IAV infection of WT (n=3) and SOD1893A(n=4) mice. **H)** Viral growth curves in WT and SOD1893A mouse embryonic fibroblasts (MEFs) infected with IAV (MOI = 3). Means and standard error of the mean (SEM) are shown. Statistics were obtained by a Student’s T-test, one-way ANOVA with Tukey post-hoc test, and Mantel-Cox test. ***, p < 0.001. ****, p < 0.0001, ns indicates not significant.

### WT and SOD1^G93A^ mice experience similar acute disease characteristics following viral infection

Given that a single acute IAV infection was associated with marked acceleration in ALS disease progression, we next sought to determine whether this was related to differences in infection kinetics and/or severity in SOD1^G93A^ mice compared to WT controls. To this end, mice were infected i.n. with a sublethal dose of IAV at day 60 of age and were subsequently monitored for influenza-associated morbidity and mortality. Both IAV-infected WT and SOD1^G93A^ mice had similar weight loss kinetics, with peak weight loss at 8 dpi (**Fig. 1E**). Likewise, no significant differences in survival following acute infection were observed (**Fig. 1F**).

Consistent with the lack of observed differences in infection kinetics and severity, there were no differences in lung viral titres between SOD1^G93A^ and WT mice as assessed by plaque forming unit (PFU) assay during peak viral load (4 dpi) (**Fig. 1G**). These findings were further confirmed using mouse embryonic fibroblasts (MEFs) derived from SOD1^G93A^ and WT mice, as infection with IAV at a multiplicity of infection (MOI) of 3 resulted in no significant differences in IAV titres in the supernatant of infected cells over the course of 48 hours (**Fig. 1H).** Taken together, these results suggest that both WT and SOD1^G93A^ mice resolve acute infection similarly, indicating that SOD1^G93A^ mice are not inherently more susceptible to IAV infection and thus, that differences in acute infection characteristics are unlikely to explain the observed acceleration in ALS disease progression.

### WT and SOD1^G93A^ mice mount similar antiviral- and pro-inflammatory responses following IAV infection

Peripheral- and neuro-inflammation are key hallmarks of ALS and related neurodegenerative disorders, correlating with disease severity in both patients and pre-clinical models, including SOD1^G93A^ mice^26,27^. Additionally, there is growing recognition that the cGAS-STING pathway, which is involved in the host antiviral response, is important for disease progression^33–35^. Given that acute viral infections induce pro-inflammatory and antiviral responses, we next sought to determine whether differences exist in such responses following IAV infection in WT and SOD1^G93A^ mice. To this end, WT and SOD1^G93A^ MEFs were stimulated with an MOI of 10 of UV-inactivated IAV (**Fig. 2A**). Viral inactivation was performed to eliminate antiviral gene antagonism by virally-encoded proteins which would confound measurement of the host antiviral response^36^. RNA was isolated from cellular lysates collected at 4-, 8-, and 24-hours post-infection for RT-qPCR to determine relative gene expression values for bona fide antiviral genes, including interferon-induced transmembrane protein 3 (*IFITM3*), RIG-I (*DDX58*), and interferon-induced protein with tetratricopeptide repeats 1 (*IFIT1*). Relative expression levels were determined using the 2^-ΔΔCt^ method and values were normalized to untreated, uninfected WT MEFs (**Fig. 2B)**. As expected, all antiviral genes were upregulated following exposure to inactivated virus in a time-dependent manner, but there were no significant differences in antiviral expression levels between the two infected groups (**Fig. 2B**).

**Figure 2.**
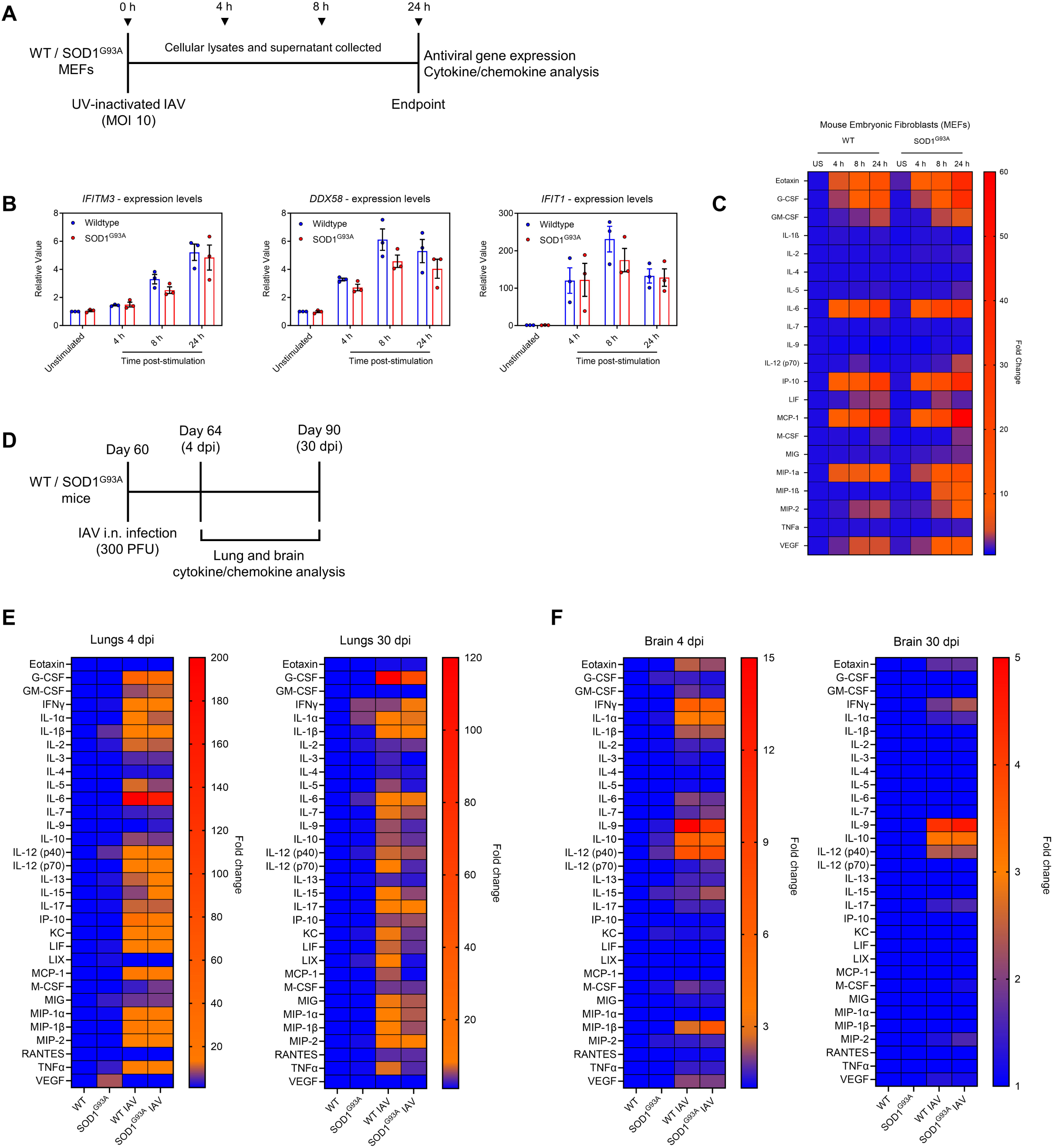
WT and SOD1G93A mice mount similar antiviral- and pro-inflammatory responses following IAV infection. **A)** Schematic of experimental timeline for assessing antiviral gene and cytokine/chemokine expression in MEFs. **B)** RT-qPCR of antiviral gene transcript levels in MEF cellular lysates following UV-inactivated IAV stimulation (MOl=10), n=3. Values are normalized to unstimulated WT MEFs C) Pro-inflammatory cytokine expression levels in the supernatant of MEFs following stimulation with UV-inactivated IAV (MOI = 10 equivalent) n=5. Values are normalized to unstimulated (US) WT MEFs. D) Experimental timeline for assessing cytokine/chemokine expression in the lungs and brain. Pro-inflammatory cytokine expression levels in E) the lungs (n=3) and F) the brain (n=3) of uninfected WT and SOD1G93A mice and !AV-infected WT and SOD1G93A mice at 4- and 30- dpi. Values are normalized to uninfected WT mice. Means and standard error of the mean (SEM) are shown.

To assess pro-inflammatory cytokines/chemokines induced by IAV infection, a 32-plex pro-inflammatory cytokine/chemokine array was performed on the supernatants collected following UV-inactivated IAV stimulation of WT and SOD1^G93A^ MEFs (**Fig. 2A**). Compared to baseline untreated MEFs, both WT and SOD1^G93A^ MEFs produced elevated levels of inflammatory analytes at all timepoints, with expression levels increasing throughout the duration of the stimulation (**Fig. 2C**). Once again, we observed no significant differences in the magnitude of cytokine/chemokine examined at each timepoint for IAV-stimulated WT and SOD1^G93A^ MEFs. To determine whether these similarities persisted *in vivo*, WT and SOD1^G93A^ mice were infected i.n. with a sublethal dose of IAV at day 60 of age and lung and brain homogenates were collected at 4- and 30- dpi (**Fig. 2D**). As expected, multiple pro-inflammatory analytes were similarly elevated in the lungs (**Fig. 2E, left panel**) and brain (**Fig. 2F, left panel**) of WT and SOD1^G93A^ mice following IAV infection at 4 dpi, as compared to uninfected mice. Strikingly, several of these analytes remained elevated at 30 dpi in the lung (**Fig. 2E, right panel**) and brain (**Fig. 2F, right panel**), well after resolution of IAV infection (**Supplemental Fig. 1**). In concordance with the observations in MEFs, there were no significant differences in pro-inflammatory cytokine/chemokine relative expression levels between IAV-infected WT and SOD1^G93A^ mice. Taken together, these data suggest that WT and SOD1^G93A^ mice mount similar antiviral and inflammatory responses to IAV infection – results that are in alignment with the observed similarities weight loss and viral replication between the two genotypes.

### Virus-induced systemic and neuroinflammation is associated with accelerated ALS disease progression

Having established that IAV infection accelerates ALS disease progression in SOD1^G93A^ mice, we next went on to determine the contribution of the immune response stimulated by the virus on disease acceleration. By inactivating the virus prior to inoculation, the contribution of viral replication to observed phenotypes is eliminated, and any differences can be attributed primarily to the host response resulting from viral detection. To this end, we administered 50 μg of formalin-inactivated IAV intramuscularly (i.m.) within the quadricep muscle and compared these to mice receiving i.m. vehicle control (equimolar formalin diluted in PBS) (**Fig. 3A**). We observed no weight loss following inoculation during the 2-week monitoring period (data not shown). Mice were then subjected to the rotarod test and monitored until endpoint was reached due to progression of ALS clinical signs. SOD1^G93A^ mice administered formalin-inactivated IAV i.m. performed significantly worse on the rotarod test, reaching 50% of their initial fall time significantly faster than mice administered i.m. vehicle control (**Fig. 3B**). Additionally, this route of administration significantly shortened lifespan of SOD1^G93A^ mice compared to those administered vehicle control (**Fig. 3C**). Since IAV infection results in considerable systemic inflammation, and the hindlimbs where inactivated IAV was injected is the site where ALS disease manifests in the SOD1^G93A^ mouse model, we set out to determine whether local inflammation in tissues other than the quadriceps also accelerated disease. To this end, we administered inactivated IAV or vehicle control intranasally (**Fig. 3A**). Similar to inactivated IAV administered i.m., no weight loss was observed (data not shown). However, in contrast to inactivated IAV given i.m., when an equal dose of formalin-inactivated IAV was administered i.n., there were no significant differences in rotarod performance (**Fig. 3D**) and there were no differences in survival due to ALS clinical signs for mice administered inactivated IAV i.n. compared to mice administered vehicle control (**Fig. 3E**).

**Figure 3.**
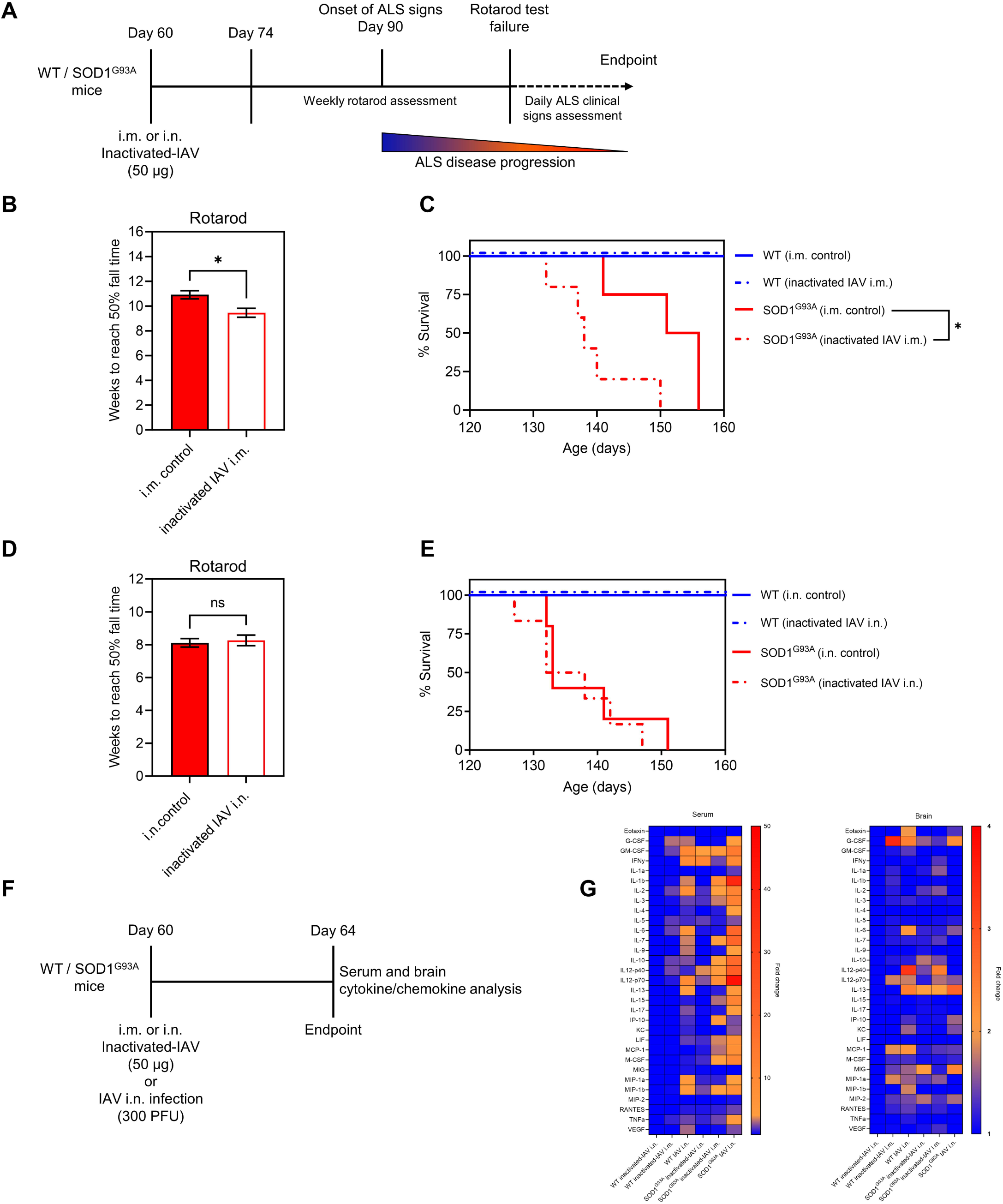
Virus-induced systemic and neuroinflammation is associated with accelerated ALS disease progression. **A)** Schematic of experimental timeline. **B)** Weeks for mock-injected WT (n=4) and SOD1G93A (n=4) and i.m. inactivated-lAV treated (50 µg) WT **(n=11)** and SOD1G93A (n=5) mice to reach 50% of their initial fall time. **C)** ALS-related endpoint of WT and SOD1G93A mice administered inactivated-lAV i.m. compared to mice given vehicle control i.m. **D)** Weeks for mock-inoculated WT (n=6) and SOD1G93A (n=5) and i.n. inactivated-lAV treated (50 µg) WT (n=8) and SOD1G93A (n=6) mice to reach 50% of their initial fall time. **E)** ALS-related endpoint of WT and SOD1G93A mice administered inactivated-lAV i.n. compared to mice given vehicle control i.n. **F)** Schematic of experimental timeline to assess cytokine/chemokine production. **G)** Heat-map of cytokine/chemokine expression levels in the serum and brain 4 days post-administration. Means and standard error of the mean (SEM) are shown. Statistics were obtained by a Student’s T-test and Mantel-Cox test. *, p < 0.05, ns indicates not significant.

To measure the magnitude of systemic and neuroinflammation induced by administering inactivated IAV relative to a bona fide IAV infection, we administered 50 μg inactivated IAV either i.n. or i.m. and compared these mice to those given 300 PFU i.n. of live IAV. At 4 days post-administration, serum and brain homogenates were obtained and a 32-plex cytokine/chemokine analysis was conducted (**Fig. 3F**). In serum, we observed the highest magnitude of pro-inflammatory analytes in mice infected with 300 PFU IAV. Additionally, an intermediate phenotype was observed in SOD1^G93A^ mice receiving inactivated IAV i.m., as levels were higher than mice administered inactivated IAV i.n. (**Fig. 3G**). The more pronounced inflammatory responses induced by live virus infection and i.m. administration of inactivated virus were therefore associated with disease progression relative to i.n. administration of inactivated IAV, which induced minimal systemic and neuroinflammation and did not accelerate disease.

Collectively, these data suggest infection-induced immunity appears to be an important factor in eliciting ALS disease acceleration. This is especially evident when inflammation is systemic or localized within disease susceptible tissues, such as the hind-limbs of SOD1^G93A^ mice.

### Acute IAV infection is associated with elevated gliosis and ALS-related transcripts in the lumbar spine

Neuroinflammation and pathological non-cell autonomous processes such as aberrant glial cell activation are critical drivers of motor neuron degeneration in patients and pre-clinical models^37^. In the SOD1^G93A^ model, selective ablation of this mutant gene in microglia and astrocytes leads to slower disease progression, suggesting a key role for these cells in ALS pathophysiology^38,39^. Furthermore, it has been well established that neuroinflammation can prime microglia to a predominantly pathogenic cytotoxic/pro-inflammatory M1 phenotype^40^. Given these findings, and that acute infection with a non-neurotropic strain of IAV was associated with sustained increased levels of pro-inflammatory cytokines/chemokines in the brain (**Fig. 2F**), we next investigated whether IAV infection induced inflammation and/or gliosis in the lumbar spine of SOD1^G93A^ mice.

To characterize the inflammatory profile, a 32-plex pro-inflammatory cytokine/chemokine analysis was performed on lumbar spine homogenates at 4 dpi (during acute infection) and day 120 of age (60 dpi – late-stage ALS disease). A trend towards increased secretion of eotaxin, IL-6, IL-10, and interferon gamma produced protein-10 (IP-10) was observed in IAV infected SOD1^G93A^ mice compared to uninfected mice at 4 dpi (**Supplemental Fig. 2A**). Next, we went on to assess whether inflammatory cytokine/chemokine expression levels following IAV infection were sustained even after the resolution of acute infection. We observed low levels of pro-inflammatory cytokines in the lumbar spine of WT mice for all analytes assessed at day 120 of age (60 dpi), regardless of previous IAV infection. SOD1^G93A^ mice had higher expression levels of multiple analytes, however, we observed no significant differences between previously IAV-infected and uninfected SOD1^G93A^ mice at this timepoint (**Supplemental Fig. 2B**), suggesting that inflammatory cytokine/chemokine levels normalize to uninfected levels by 60 dpi.

To assess the level of gliosis in the lumbar spine, WT and SOD1^G93A^ mice were infected i.n. with a sublethal dose of IAV at day 60 of age and immunohistochemistry was performed at day 90 of age (30 dpi) (**Fig. 4A**). This timepoint was chosen as sustained peripheral and central inflammation following sublethal IAV infection was observed (**Fig 2E and 2F**), and that day 90 typically corresponds to onset of ALS clinical signs in this model. Spinal sections were isolated and fixed sections were probed with anti-GFAP and anti-Iba1 antibodies to measure levels of astrocyte and microglia activation, respectively. In agreement with previously published studies, elevated levels of gliosis as measured by the number of GFAP+ astrocytes (**Fig. 4B**) and Iba1+ microglia (**Fig. 4C**) were observed in lumbar spines from uninfected SOD1^G93A^ mice in comparison to uninfected WT controls. Strikingly, whereas there we no significant increases in gliosis in WT mice following IAV infection, this single infection event 30 days earlier was associated with both a marked increase in GFAP+ astrocytes, and a significant increase in Iba1+ microglia in the lumbar spine of SOD1^G93A^ mice. Furthermore, gliosis in the SOD1^G93A^ mice was associated with a trend towards fewer motor neurons in the lumbar spine at day 90 of age (30 dpi), as measured by ChAT+ cells (**Supplemental Fig. 3**).

**Figure 4.**
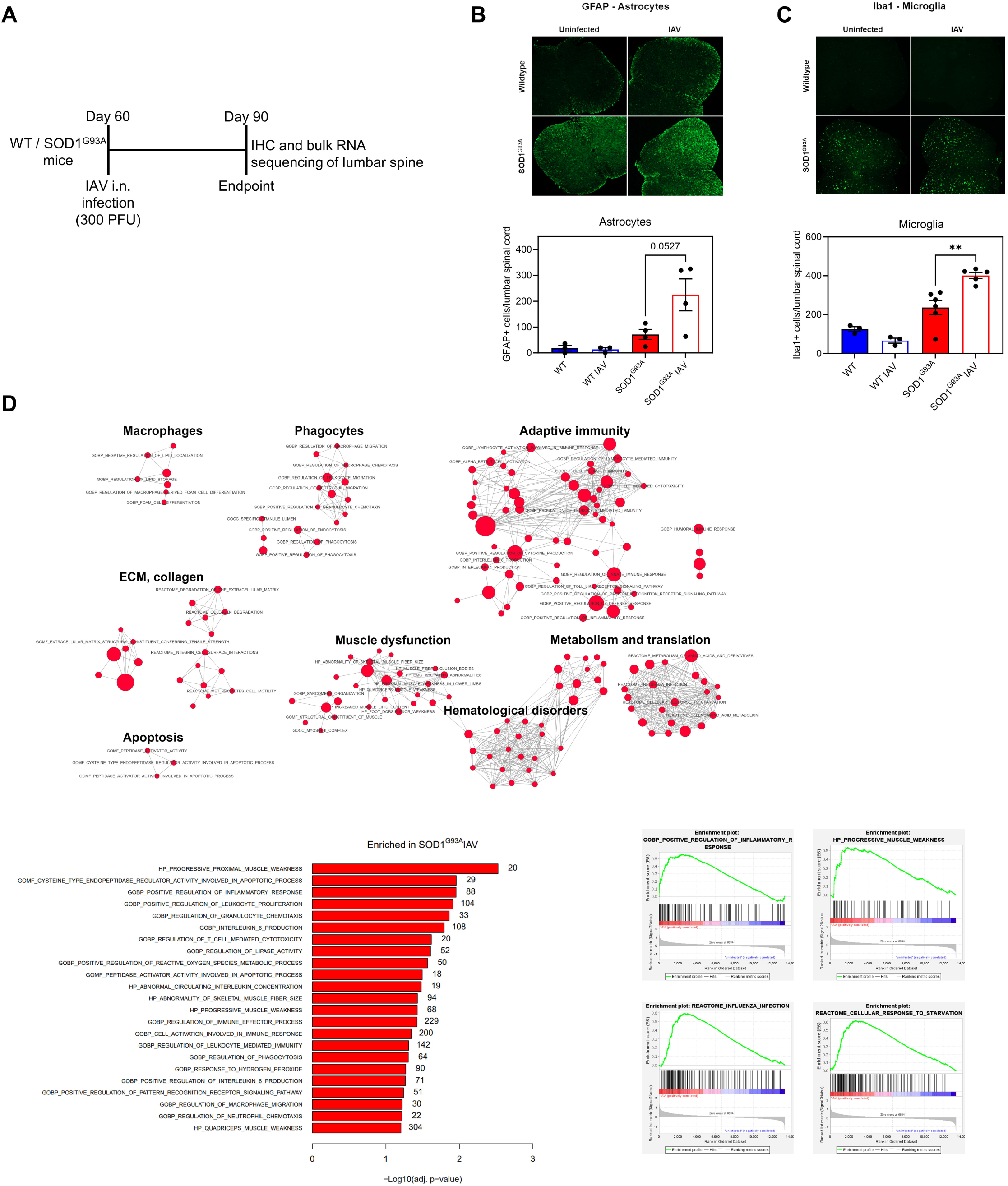
Acute IAV infection is associated with elevated gliosis and ALS-related transcripts in the lumbar spine. **A)** Schematic for assessing glial cell activation and transcriptomics in the lumbar spine. **B)** Representative images of the lumbar spine sections probed for GFAP **(top)** and quantification of GFAP+ cells for each lumbar spine at day 90 of age (30 dpi) **(bottom). C)** Representative images of the lumbar spine sections probed for lba1 **(top)** and quantification of lba1+ cells for each lumbar spine at day 90 of age (30 dpi) **(bottom).** Images were taken at 5X magnification. **D)** Gene set enrichment analysis of pathways significantly enriched in IAV infected SOD1G93Amice (n=3) compared to uninfected SOD1G93Amice (n=3) at day 90 of age (30 dpi). Enrichment map **(top),** select enriched pathways with the number of genes associated with each pathway listed beside the bars **(bottom left),** individual enrichment score plots **(bottom right).** Means and standard error of the mean (SEM) are shown. Statistics were obtained by a one-way ANOVA with Tukey post-hoc test. **, p < 0.01.

Additionally, RNA was isolated from the lumbar spine of both previously IAV infected and uninfected SOD1^G93A^ mice at 90 days of age (30 dpi) and subjected to RNA sequencing to identify differentially expressed genes. Gene set enrichment analysis (GSEA) revealed significant upregulation in pathways associated with innate and adaptive immunity, lipid metabolism, apoptotic pathways, and muscle dysfunction in previously IAV infected SOD1^G93A^ mice compared to uninfected SOD1^G93A^ mice (**Fig 4D**). Specifically, pathways related to inflammatory and immune responses were prominently elevated, indicating a robust immune activation in the lumbar spine well after IAV infection was resolved. Overall, these results suggest that prior acute viral infection potentiates gliosis in the lumbar spine and induction of transcriptional programs relating to inflammation, immunity, and lipid and muscular dysfunction.

### Inhibition of microglial activation during acute IAV infection protects SOD1^G93A^ mice from accelerated disease progression

We have thus far demonstrated that a single acute infection event prior to onset of ALS clinical signs accelerates ALS disease progression in SOD1^G93A^ mice, and resulted in elevated inflammatory signatures and gliosis in the lumbar spine. Consequently, we next sought to determine whether inhibiting this activation during acute infection could prevent ALS disease acceleration. Minocycline is known to have anti-inflammatory properties and inhibit aberrant microglial activation^41^. Importantly, minocycline has been shown to improve ALS outcomes in transgenic SOD1 mice by inhibiting microglial activation^42^. To test whether inhibiting microglia activation during acute infection prevents IAV-induced ALS acceleration, SOD1^G93A^ mice were administered 50 mg/kg minocycline via intraperitoneal (i.p.) injection daily. Treatment was initiated one day prior to infection (day 59 of age) and continued until 3 weeks post-infection (day 81 of age) (**Fig. 5A**). Both minocycline treated and untreated mice infected with IAV experienced similar kinetics of weight loss following infection, with weight loss peaking around 8 dpi (**Fig. 5B**), indicating that treatment did not alter acute disease kinetics in comparison to controls. At 2 weeks post-infection, mice were assessed on the rotarod. SOD1^G93A^ mice administered minocycline performed significantly better on the rotarod test with a significant delay in minocycline treated mice reaching 50% of their initial fall time compared to untreated mice infected with IAV (**Fig. 5C**). However, there were no observable differences in survival due to ALS clinical sign progression between the two groups. The shape of the Kaplan-Meyer curve alongside the rotarod data suggest that the benefits of minocycline may diminish upon cessation of treatment (**Fig. 5D**). We next assessed levels of microglial activation in the lumbar spine at the time of treatment cessation (day 81 of age – 3 weeks post-infection). Indeed, we observed a trend towards decreased Iba1+ cells in minocycline treated SOD1^G93A^ mice infected with IAV compared to those infected receiving vehicle control (**Fig. 5E**). Altogether, these results indicate that inhibiting microglial activation during acute infection helps to prevent infection-induced disease acceleration.

**Figure 5.**
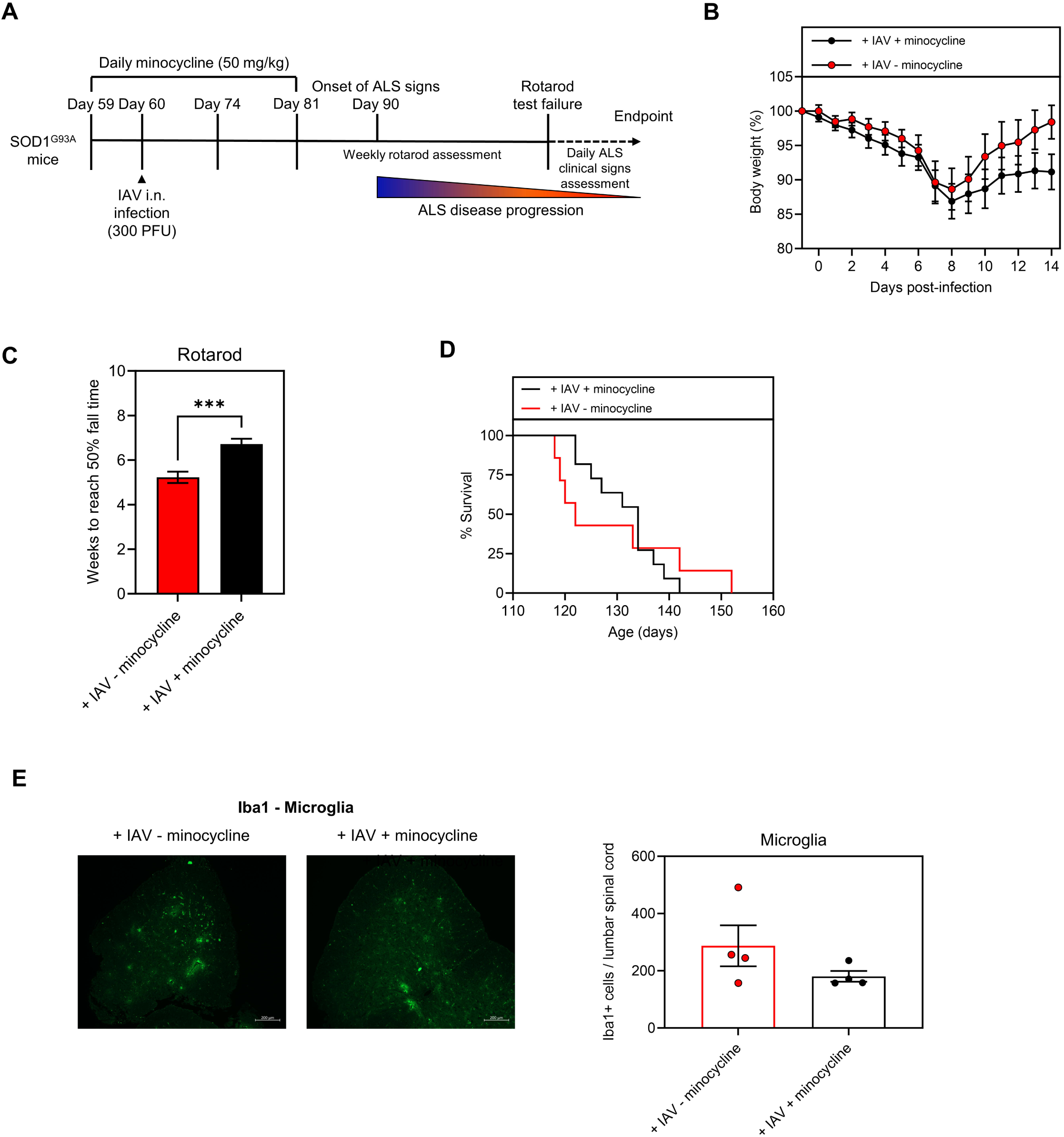
Inhibition of microglial activation during acute IAV infection protects S0D1G93A mice from accelerated disease progression. **A)** Schematic of experimental timeline. SOD1893Amice were administered 50 mg/kg minocycline daily at day 59 of age (one day prior to IAV infection) until day 81 of age (3 weeks post-infection). **B)** Weight loss of minocycline treated (n=12) and control (n=8) SOD1893Amice following IAV infection. **C)** Weeks for minocycline treated and IAV infected (n=11) SOD1893Amice and IAV infected SOD1893Amice given vehicle control (n=7) to reach 50% of their initial fall time. **D)** Survival of IAV infected minocycline treated mice compared to IAV infected control. **E)** Representative images **(left)** and lba1 quantification **(right)** in the lumbar spine at day 81 of age (administration cessation) of IAV infected SOD1893Amice given minocycline or vehicle control. Images were obtained at 5X magnification. Means and standard error of the mean (SEM) are shown. Statistics were obtained by a Student’s T-test, and Mantel-Cox test. ***, p < 0.001.

### Oseltamivir treatment during IAV infection protects SOD1^G93A^ mice from accelerated disease progression

We have shown that prior IAV infection potentiates gliosis in lumbar spine of SOD1^G93A^ mice, and that administering minocycline reduces IAV-induced glial activation resulting in subsequent motor function improvements. We next examined the protective role of administering a direct-acting antiviral on infection-induced ALS disease acceleration. Oseltamivir is a direct-acting antiviral that prevents influenza illness by inhibiting the activity of neuraminidase on influenza viruses^43^. To this end, SOD1^G93A^ mice were infected i.n. with a sublethal dose of IAV at day 60 of age and were subsequently administered 10 mg/kg oseltamivir phosphate via oral gavage twice daily. Oseltamivir administration was initiated 1 dpi and continued for 7 consecutive days (**Fig. 6A**). As expected, oseltamivir administration attenuated disease, reducing weight loss following IAV infection compared to mice administered vehicle control (**Fig. 6B**). Additionally, IAV-infected mice administered oseltamivir had better motor performance on the rotarod test compared to infected mice that were administered vehicle control (**Fig. 6C**). However, similar to minocycline treatment, this did not result in a significant lengthening of survival, though there was a positive trend (**Fig. 6D**). Collectively, these data demonstrate the therapeutic benefit of antiviral treatment in preventing accelerated ALS motor decline resulting from IAV infection.

**Figure 6.**
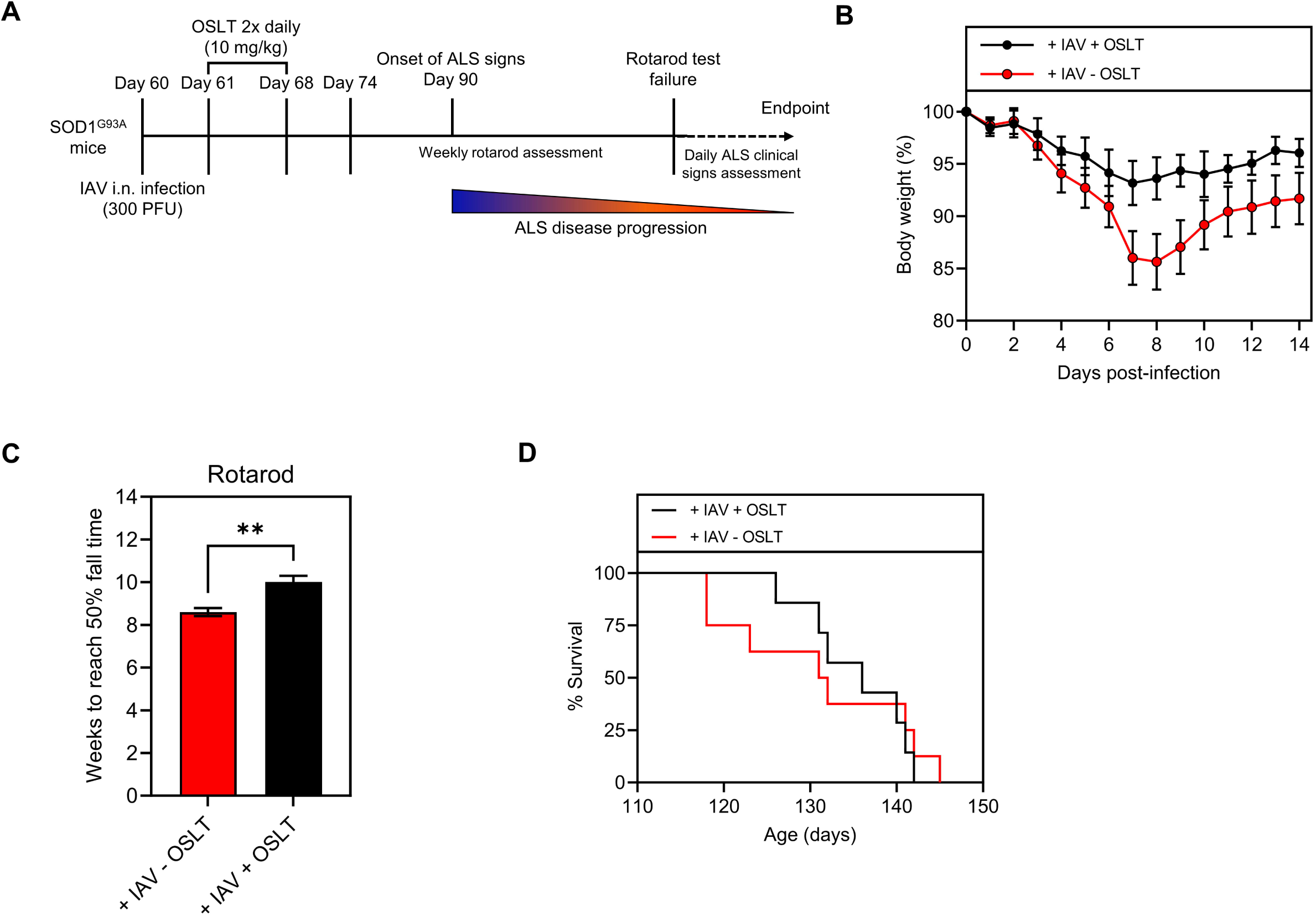
Oseltamivir treatment during IAV infection protects SOD1G93A mice from accelerated disease progression. **A)** Schematic of experimental timeline. SOD1G93A mice were infected with 300 PFU IAV and treated with oseltamivir (OSLT) via oral gavage twice daily at a dose of 10 mg/kg 1 day following infection. **B)** Weight loss of IAV infected OSLT treated (n=6), IAV infected untreated (n=9) SOD1893A mice. **C)** Weeks for IAV infected OSLT treated (n=7) and IAV infected vehicle control treated (n=7) mice to reach 50% of their initial fall. **D)** Survival of OSLT treated and untreated, IAV infected SOD1893A mice due to ALS clinical signs. Means and standard error of the mean (SEM) are shown. Statistics were obtained by Student’s T-test, and Mantel-Cox test. **, p < 0.01.

### SARS-CoV-2 infection prior to ALS onset accelerates disease progression in SOD1^G93A^ mice

Given the widespread global impact of COVID-19, we next sought to determine whether similar ALS disease acceleration was observed in SOD1^G93A^ mice acutely infected with SARS-CoV-2. To this end, SOD1^G93A^ mice were infected i.n. with a sublethal dose (10^5^ PFU) of a mouse-adapted SARS-CoV-2 (strain MA10) (**Fig. 7A**). We observed that SOD1^G93A^ mice infected with SARS-CoV-2 had worse motor performance assessed through the inverted grip test (**Fig. 7B**). However, in contrast to acute IAV infection, acute SARS-CoV-2 infection did not accelerate time to endpoint due to ALS clinical signs faster than uninfected controls (**Fig. 7C**). Next to assess SARS-CoV-2 infection disease kinetics in this model, SOD1^G93A^ mice were infected i.n. with a sublethal dose of SARS-CoV-2. We observed no appreciable weight loss throughout the course of infection (**Fig. 7D**), and in concordance with IAV infection, WT and SOD1^G93A^ mice experience similar viral burden within the lungs (**Fig. 7E**). Taken together, these results suggest that a sublethal SARS-CoV-2 infection during the pre-symptomatic stages of ALS significantly accelerates motor decline.

**Figure 7.**
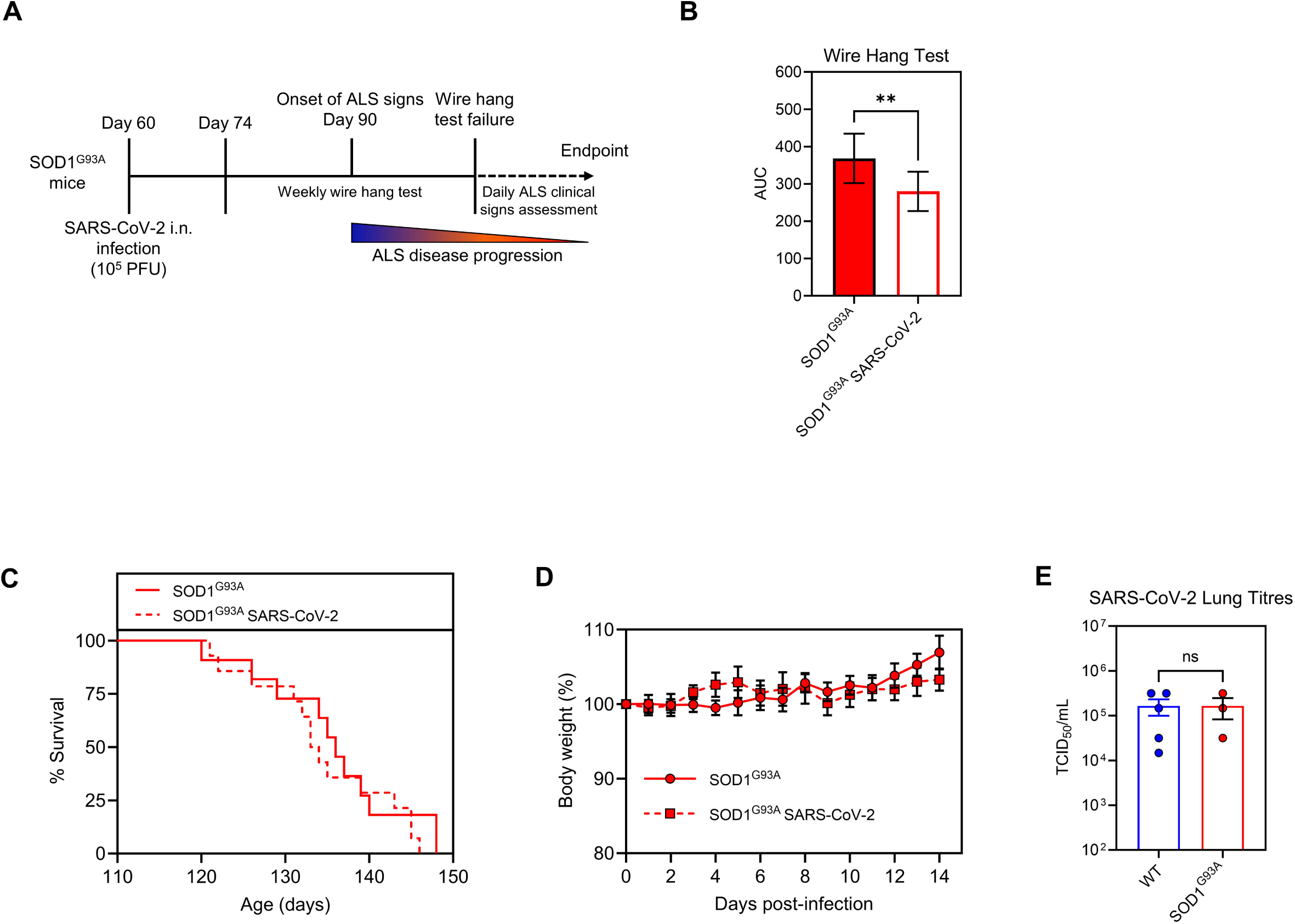
SARS-CoV-2 infection prior to ALS onset accelerates disease progression in S0D1893A mice. **A)** Schematic of the experimental timeline for SARS-CoV-2 infection of mice. **B)** Wire hang test evaluation (area under the curve (AUC)) of uninfected SOD1893A(n = 11) and SARS-CoV-2 infected SOD1893A(n = 12) mice. **C)** Survival of uninfected SOD1893A (n = 11) and SARS-CoV-2 infected SOD1893A(n = 12) mice due to ALS-related endpoint. **D)** Weight loss following SARS-CoV-2 infection, n = 5 - 6. **E)** Viral titres in the lungs at 2 dpi (n = 4). AUC analysis depicts standard deviation. Statistics were obtained by a Student’s T-test and Mantel-Cox test.**, p < 0.01, ns indicates not significant.

## Discussion

ALS remains an incurable disease and current treatments have relatively modest therapeutic benefits. Tremendous efforts have been made to develop new treatments, however many clinical trials have eventually failed to slow disease progression as measured by the ALSFRS-R^44,45^. The immense heterogeneity of ALS is an important factor contributing to this lack of success, a product of genetic and environmental factors^46^. Viral infections have long been associated with ALS and other neurodegenerative diseases. However, most prior studies have been based on observational studies that are correlative in nature. Additionally, these studies have largely focused on chronic and neurotropic infections, as these are more easily detected in post-mortem and serological analyses of ALS patients. Indeed, infections that occur and are resolved throughout the life of an individual can have profound long-term impacts (e.g. “long COVID”).

Importantly, the mechanism underlying viral-mediated potentiation of ALS disease progression has been widely understudied. Our study demonstrates that a single, acute infection with common pathogens, such as IAV and SARS-CoV-2, during the pre-symptomatic stages of disease accelerates the progression of ALS in a murine model. Our work also suggests that potentiation of aberrant inflammatory immune processes in the CNS may be a unifying mechanism to explain why so many viruses with diverse biologies have been implicated in ALS.

Here, we used the SOD1^G93A^ model to explore the mechanisms through which viral infections impact ALS disease. This is one of the most extensively characterized mouse models of ALS with tractable disease features that closely resemble “classical” ALS^47,48^. Additionally, timing of disease onset and rate of progression are predictable and have been widely documented, allowing us to investigate whether infection during pre-symptomatic stages has an influence on disease progression^48^. A mouse-adapted strain of IAV subtype H1N1 (PR8) and mouse-adapted SARS-CoV-2 (strain MA10) were used as model pathogens due to their ubiquity. Importantly, previous studies indicate that these particular isolates are not neuroinvasive, yet infection induces signatures of neuroinflammation through peripheral perturbations^19,49,50^.

We found that a sublethal i.n. infection with IAV at day 60 of age resulted in poorer motor function and significant acceleration of disease relative to uninfected SOD1^G93A^ mice. Disease acceleration was also observed in SOD1^G93A^ mice infected with SARS-CoV-2. This acceleration occurred despite clearance of the virus, as no detectable virus was found in the lungs or brains of infected WT and SOD1^G93A^ mice following infection resolution at 14 dpi^19^. Furthermore, we observed no differences in the dynamics of acute infection experienced by WT and SOD1^G93A^ mice following IAV and SARS-CoV-2, with both genotypes demonstrated similar weight loss trajectories and viral burden in the lungs following infection. This suggests that individuals at risk of developing ALS may not be inherently more susceptible to infection prior to onset of clinical disease.

Neuroinflammation is a well-established feature of ALS and other related neurodegenerative diseases, like Parkinson’s Disease and Alzheimer’s Disease. Indeed, as we and others have shown, many genes associated with ALS have important roles in regulating innate and adaptive immune responses to infection^51–53^. ALS disease progression is correlated with elevated levels of soluble pro-inflammatory mediators in the peripheral blood and CSF, and there is evidence of substantial cross-talk between the periphery and CNS^26,27^. Interestingly, we observed elevated levels of pro-inflammatory cytokines/chemokines in the lungs and brains of infected WT and SOD1^G93A^ mice at 4 dpi that remained elevated 30 dpi, well after the initial infection had been resolved. This suggests long-term neuroinflammation can be caused by acute infection originating from peripheral responses. To distinguish the relative contribution of infection-induced immunity on ALS acceleration relative to that of virus replication itself, inactivated IAV was administered through i.m. or i.n. inoculation. We observed that i.m. administration in the quadriceps resulted in accelerated ALS disease progression of SOD1^G93A^ mice relative to those administered vehicle control. However, this was not observed when inactivated IAV was i.n. administered. Taken together, our findings show that inflammation induced by viral exposure (and not exclusively the direct effects of viral replication alone), are important for accelerating ALS disease progression. Additionally, the location in which inflammation is generated is critical. Systemic inflammation and inflammation generated in disease susceptible tissue, such as the hindlimbs in the SOD1^G93A^ model, appears to be especially deleterious to these mice, whereas inflammation localized in tissues distal from the sites of disease, such as the lungs, did not accelerate progression.

To gain further mechanistic insight, immunohistochemical analysis of the lumbar spine provided evidence of elevated astrocyte and microglial activation levels in previously infected SOD1^G93A^ mice compared to control mice at day 90 of age (30 dpi). Given the reported importance of glial cells in ALS pathophysiology, the observation that acute infection results in sustained gliosis, long after the infection is cleared, provides a compelling mechanism through which these infections potentiate ALS. The immunohistochemical findings were further corroborated through bulk RNA sequencing of the lumbar spinal cord of previously IAV infected and control SOD1^G93A^ mice. GSEA revealed significantly enriched transcripts for pathways relating to inflammation, immunity, lipid and muscle dysregulation and apoptosis in IAV infected SOD1^G93A^ mice. As previously mentioned, soluble pro-inflammatory mediators and the induction of innate and adaptive immunity are positively correlated with ALS progression. However, interestingly, enrichment of lipid dysregulation and apoptotic pathways suggests additional changes in bioenergetics and intrinsic cellular processes in the lumbar spine following IAV infection. Indeed, dysregulated lipid profiles and apoptotic signaling have been observed in ALS patients^54–56^. Notably, gene sets related to muscle dysfunction and weakness were upregulated in IAV infected SOD1^G93A^ mice, indicating exacerbated neuromuscular impairment characteristic of this ALS model. Muscle weakness and atrophy are primary symptoms of ALS, and their exacerbation following viral infection suggests a potential interaction between acute peripheral IAV infection and neuromuscular health^57^.

Given our findings that acute viral infection accelerates disease, presumably via potentiation of pathogenic immune responses in the CNS, we sought to determine the therapeutic potential of two separate approaches: chemically inhibiting microglial activation, and treating acute viral infection using antivirals. We found that minocycline and OSLT treatment improved performance of SOD1^G93A^ mice on the rotarod. This suggests that acute treatment is efficacious but must likely be sustained following infection in order to extend survival (since we observed the resumption of accelerated disease upon treatment cessation). Indeed, prior studies showing efficacy of antivirals and immunomodulatory drugs in pre-clinical models have maintained treatment for the duration of the disease^11,58–60^.

The data presented here demonstrates that a single sublethal infection with common, acute viruses exacerbate ALS disease. These results also underscore inherent difficulties in conducting these types of studies in humans. In our model, were no observable differences in susceptibility to, or severity of acute viral infection when comparing WT and SOD1^G93A^ mice. Furthermore, acceleration of disease was not evident until several weeks after the acute infections had been cleared. These types of “hit-and-run” mechanisms are inherently challenging to study in epidemiological contexts – especially for ubiquitous pathogens wherein differences in seroprevalence would not be expected. Elegant recent work using data from European biobanks, highlighted extensive associations between infections and many neurodegenerative diseases, including ALS, even 15 years following an exposure^17^. Here, we build on those studies and provide mechanistic insight that may explain how and why pathogens with inherently different biologies all contribute to neurodegenerative disease – namely, through their shared propensity to elicit inflammatory immune responses. To our knowledge, this is the first study investigating whether a direct causal relationship exists with acute viral infections (particularly non-neurotropic infections), and ALS progression. This should set the stage for additional studies in human cohorts at risk for developing ALS.

### Limitations

For practical reasons, our studies were limited to the SOD1^G93A^ mouse model, which is one of the few ALS models that recapitulates key features of neuromuscular disease with predictable and tractable kinetics. In the future, it would be interesting to validate that our findings extend to other ALS models with distinct genetic backgrounds (most of which do not show a neuromuscular phenotype, but may demonstrate features consistent for frontotemporal dementia, for example). Likewise, it would be interesting to determine whether infection could serve as a trigger for ALS in a model wherein animals do not spontaneously develop disease. However, a suitable “inducible” model of ALS has yet to be developed, to our knowledge.

## Materials and Methods

### Animal colony and ethics statement

All mouse experiments were approved by the McMaster University Animal Research Ethics Board. SOD1 Tg mice expressing the G93A mutation, raised on the B6SJL-TgN(SOD1*G93A)1Gur background (SOD1^G93A^), and wildtype BL6SJL (WT) mice were purchased from the Jackson Laboratory. Mice were maintained on a 12 hour light/dark cycle and low-fat dietary chow and water was provided *ad libitum*. Breeding pairs consisted of male SOD1^G93A^ mice and female WT mice, and mice were bred up to the second generation. Litters were weaned at 21 days of age and housed at a maximum of five mice per cage. All experimental mice were age-matched.

### Viruses and Cells

Influenza A/Puerto Rico/8/1934/H1N1 (PR8) was propagated in 10 day-old specific-pathogen-free embryonated chicken eggs (Canadian Food Inspection Agency)^61^. Viral stock concentrations were quantified by performing plaque assay on Madin-Darby Canine Kidney (MDCK) cells (ATCC, VA, USA). SARS-CoV-2 MA10 was kindly provided by Dr. Ralph Baric^62^. MDCKs and VeroE6 cells (ATCC, VA, USA) were maintained in Dulbecco’s minimal essential media (DMEM) (Gibco) supplemented with 1 X GlutaMAX^TM^ supplement (Gibco), 10% heat inactivated fetal bovine serum (FBS) (Gibco), 100 U/mL penicillin and 100 µg/mL streptomycin (Gibco), and 1% of 1 M HEPES (ThermoFisher) at 37°C and 5% CO_2_. Mouse embryonic fibroblasts (MEFs) were isolated from embryos 13.5 days post-conception as previously described^63^. MEFs were maintained in DMEM supplemented with 1 X GlutaMAX^TM^ supplement, 15% FBS, and 100 U/mL penicillin and 100 µg/mL streptomycin and incubated at 37°C and 5% CO_2_.

### IAV and SARS-CoV-2 Infection

WT and SOD1^G93A^ mice were anesthetized using isoflurane and infected with IAV i.n. in a 40 µL inoculum of PBS. Mice were infected with 300 plaque forming units (PFU) of IAV, equating to a 0.1 median lethal dose (LD_50_) infection. SARS-CoV-2 infections were performed using a mouse-adapted strain of SARS-CoV-2 MA10. Mice were anesthetized using isoflurane and infected with SARS-CoV-2 MA10 i.n. at a dose of 10^5^ PFU. Weight loss, lethargy, and general body condition were monitored following infection for up to 2 weeks post-infection. Endpoint due to IAV- and SARS-CoV-2-associated mortality was defined as mice reaching 80% of their pre-infection weight.

### Cytokine and Chemokine Analysis

Evaluation of cytokine and chemokine induction in the lungs, brain, and lumbar spinal cord were performed through a mouse 32-plex cytokine/chemokine array (MD32) (Eve Technologies, Alberta, Canada).

### IAV Inactivation

100 X inactivation solution (0.925% formaldehyde in PBS) (Anachemia) was combined with 4.2 mg/mL of purified IAV suspension to a final concentration of 1 X inactivation solution. Samples were incubated at 4°C on constant rotation for 72 hours, followed by storage at -80°C.

### Inactivated IAV Administration

WT and SOD1^G93A^ mice were administered 50 µg of formaldehyde-inactivated IAV in a 100 µL inoculum via i.m. injection into the quadricep muscle, and. i.n. administration was performed with a 40 µL inoculum of 50 µg of formaldehyde-inactivated IAV.

### Rotarod Test

Motor performance and general coordination was measured using the rotarod test (Panlab, Harvard Apparatus, Holliston, MA, USA). The rotarod was set to acceleration mode (4 rpm to 15 rpm in 60 s, or 0.2 rpm/s) for a maximum rotation speed of 15 rpm. WT and SOD1^G93A^ mice were tested for a maximum of 180 seconds and the latency for falling off the apparatus was recorded. Mice were assessed once weekly until they were unable to support themselves on the apparatus, with the average of six trials being used.

### ALS Clinical Sign Assessment and Endpoint Monitoring

At day 120 of age, mice were monitored daily until reaching endpoint due to ALS clinical signs. Mice were assessed based on a 4-point neurological scoring system. Score 0: full extension of hind-limbs when suspended by the tail, Score 1: collapse or partial collapse of the hind-limbs and/or trembling of the hind-limbs when suspended by the tail, Score 2: Toe curling during locomotion, Score 3: paralysis of one or both of the hind-limbs, Score 4: unable to right themselves within 30 s of being placed on either side. Endpoint was defined as mice reaching a neurological score of 4.

### IAV Plaque Assay

MDCK cells were seeded in a 6-well plate at a density of 6.5 x 10^5^ cells per well and incubated for 48 hours at 37 °C in 5 % CO2. Following incubation, samples were serially diluted in 1 X Minimum Essential Media (MEM), 1 X GlutaMAX^TM^ supplement, 1.5 % sodium bicarbonate, 200 mM HEPES, 1 X MEM Vitamin Solution, 1 X MEM Amino Acid Solution, 2 mg/mL penicillin/streptomycin, 7 % BSA). Cells were infected for 1 hour at 37 °C in 5 % CO_2_. Cells were subsequently overlayed with 1 % (m/v) agar bacteriological combined with one part 2 X MEM, 0.01 % DEAE-Dextran, 1 µg/mL N-p-Tosyl-L-phenylalanine chloromethyl ketone (TPCK)-treated trypsin and plates were incubated at 37 °C in 5 % CO_2_ for 48 hours. Cells were fixed with 3.7 % formaldehyde diluted in PBS for 30 min at RT and stained with crystal violet.

### IAV Growth Curves

WT and SOD1^G93A^ MEFs were infected with IAV at a multiplicity of infection (MOI) of 3. Supernatant was collected at 0, 2, 4, 8, 12, 24, and 48 hours post-infection and viral titres were assessed using plaque assays on MDCK cells. For *in vivo* studies, WT and SOD1^G93A^ mice were infected with 300 PFU of IAV i.n. and lungs were collected in serum-free DMEM, and homogenized using the Bullet Blender Gold High-Throughput Bead Mill Homogenizer (Next Advance). Viral titres from lung homogenates were determined via plaque assay on MDCK cells.

### Antiviral Gene Expression Quantification and Cytokine Analysis

WT and SOD1^G93A^ MEFs were plated and stimulated with UV-inactivated IAV at an MOI of 10 equivalent and supernatant was collected at 0, 4, 8, and 24 hours post-stimulation. For gene expression quantification, RNA from cell lysates was isolated using the RNeasy kit (Qiagen) and 1 μg of complimentary DNA was synthesized using the Maxima First Strand cDNA Synthesis kit, according to manufacturers instructions (Thermo Fisher Scientific). Gene expression levels were assessed using SensiFAST SYBR reaction kit (Bioline Reagents Ltd.) for quantitative real-time PCR (qRT-PCR). For cytokine analysis, supernatant from stimulated MEFs were collected at 0, 4, 8, and 24 h post-stimulation. For *in-vivo* studies, mice were infected with 300 PFU IAV i.n. and lung and brain homogenates were harvested in PBS containing Pierce Protease Inhibitor (Thermo Fisher Scientific) at 4- and 30- days post-infection. Cytokine levels were assessed using a 32-plex mouse cytokine/chemokine array (EVE Technologies Corp., Calgary, AB, Canada).

### Fluorescent Immunohistochemistry

Spinal cord sections were isolated using hydraulic extrusion as previously described, with slight modifications^64^. Briefly, mice were anesthetized and transcardially perfused with PBS. The spinal cord was removed and cut immediately distal to the 13^th^ rib to isolate the lumbar spine. The lumbar spinal cord was hydraulically removed from the vertebrae using 5 mL of PBS and stored in 3.7 % PFA for 72 hours. Subsequently, samples were sliced 5 µm thick and mounted onto paraffin. Sections were deparaffinized with xylene and subsequently dehydrated using decreasing ethanol concentrations of 100%, 95%, 95%, 75%, and 75%, for 5 min at each step. Slices were then rehydrated in H_2_O for 5 min. Antigen retrieval was performed with 1 X citrate buffer (Abcam) for 20 min at 95°C. Thereafter, slices were washed 3 times with PBS-T (0.1% Tween 20, Sigma Aldrich). Slices were permeabilized using 0.2 % Triton^TM^ X-100 diluted in PBS-T for 20 min at RT, and then washed 3 times with PBS-T. Following permeabilization, slices were blocked with 10% normal goat serum (Abcam) and 1% BSA diluted in PBS for 1 hour at RT. Spinal cord slices were incubated with primary antibodies overnight at 4°C. Primary antibodies included: Iba1 [1:300] (Abcam), GFAP [1:400] (Abcam), ChAT [1:100] (Abcam). Slices were then washed 3 times with PBS-T and probed with compatible AlexaFluor-conjugated secondary antibodies at a 1:1000 dilution (Invitrogen), washed 3 times, and mounted with EverBrite mounting medium (Biotium Inc.). Images were taken with the Zeiss Imager M2 microscope.

### Minocycline and Oseltamivir Treatment

Minocycline hydrochloride (Sigma-Aldrich) was dissolved in PBS and administered intraperitoneally (i.p.) once daily to SOD1^G93A^ at a dose of 50 mg/kg beginning one day prior to infection (day 59 of age) for a duration of three weeks following infection (day 81 of age). Mice were monitored for weight loss, assessed on the rotarod, and monitored until reaching endpoint due to ALS clinical signs. For therapeutic oseltamivir treatment, mice were infected at day 60 of age and treated twice daily with 10 mg/kg oseltamivir phosphate (Toronto Research Chemicals, O701000) via oral gavage for 7 consecutive days starting at 1 dpi. Mice were monitored for weight loss, assessed on the rotarod, and monitored until reaching endpoint due to ALS clinical signs.

### RNA Sequencing

Lumbar spinal cord was extracted and flash frozen in liquid nitrogen. Once frozen, samples within a cryotube were disrupted using a pestle and homogenized by pipetting with 600 µL of RLT buffer (Qiagen). Homogenized samples were then transferred to a Qiashredder spin column (Qiagen), and the flowthrough was subsequently spun in a gDNA eliminator column (Qiagen). Equal volume of 70% ethanol was added and RNA isolation was carried out as per the RNeasy instruction manual (Qiagen). Sample libraries were prepared using the NEBNext Ultra^TM^ II Directional RNA Library Prep Kit for Illumina (New England Biolabs) and cDNA libraries were sequenced using an Illumina HiSeq machine (2×50 bp sequence reads). The reads were trimmed using TrimGalore and then aligned with GRCm39 reference using STAR^65^. Next, the reads were counted by using HTSeq count^66^. Genes, showing low levels of expression were removed using EdgeR package in R, resulting in 12,503 genes^67^. Counts for these remaining genes were normalized with TMM normalization method and then transformed using voom transformation^68,69^. The limma package in R was used to identify genes differentially expressed between the infected and the uninfected groups of each genotype^70^. For differential expression analysis, corrected p-values < 0.05 were considered to be significant. Gene Set Enrichment Analysis (GSEA) was performed to compare WT and SOD1^G93A^ IAV infected to their uninfected counterparts respectively using C2 canonical pathways and C5 GO collections^71^. Enrichment with corrected p-value < 0.1 was considered to be significant. GSEA results were shown using Enrichment Maps in Cytoscape environment and barplots created in R^72^. Additional results were visualized in PCA plot using rgl package in R.

### Statistical Analysis

All data and figures were generated, and statistical analyses were performed using GraphPad Prism (GraphPad Software v.10., La Jolla, CA, USA).

## Supporting information

Graphical Abstract

## Key resources table

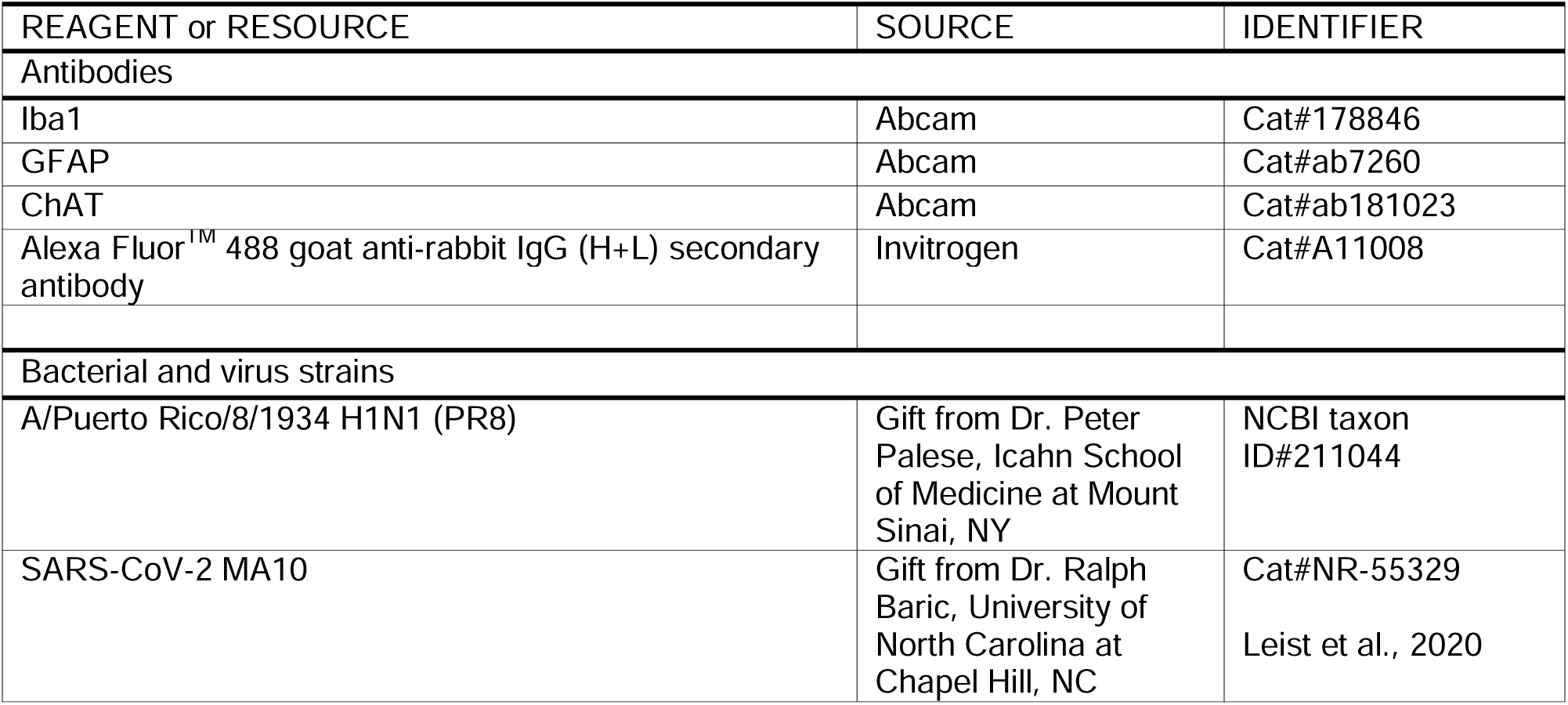
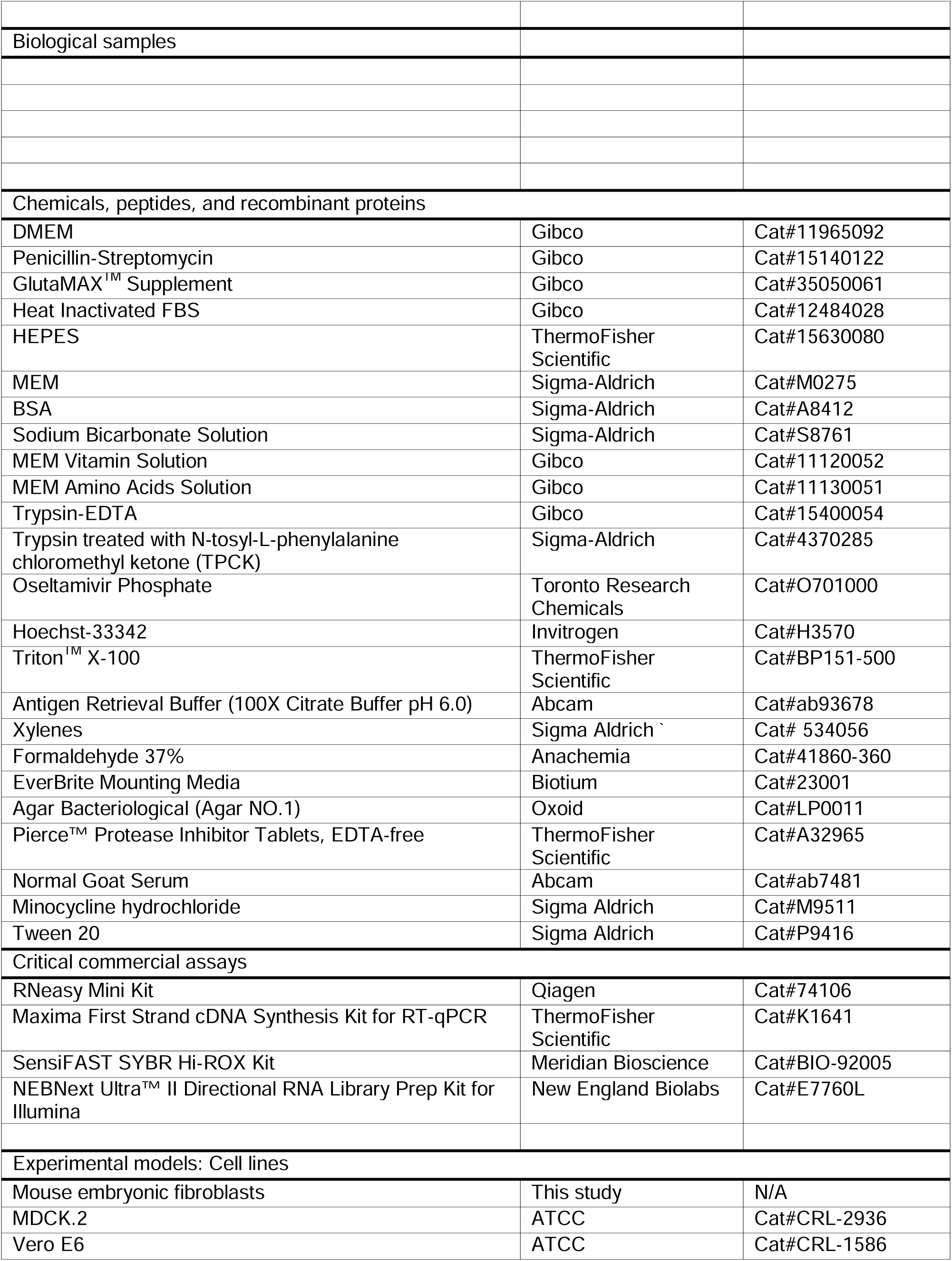
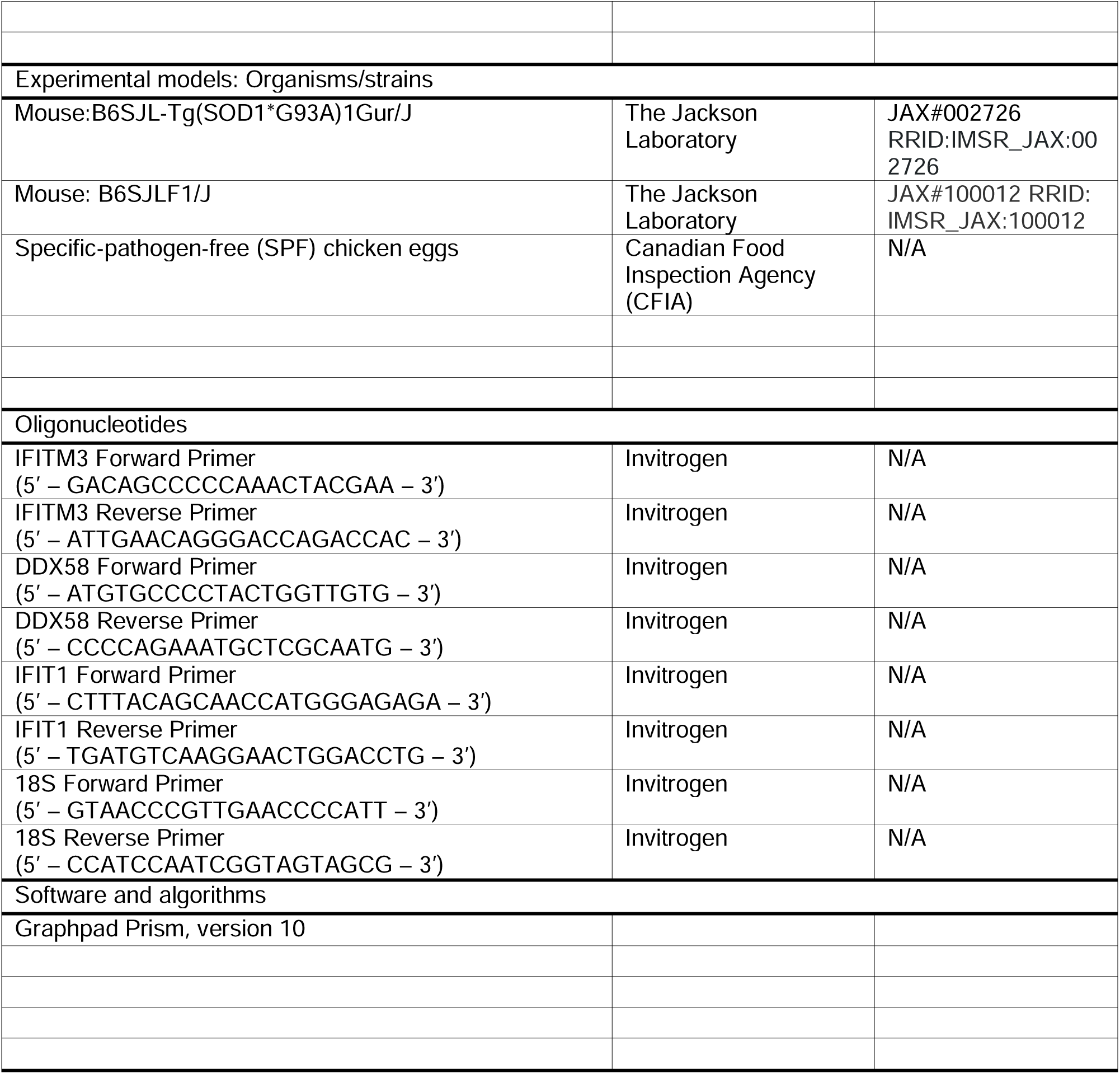

## Acknowledgement

Authors are thankful to Dr. Peter Palese for providing influenza A virus (A/Puerto Rico/8/1934 H1N1) and to Dr. Ralph Baric for providing mouse-adapted SARS-CoV-2 (strain MA10). Authors are also thankful to the McMaster Histology Core Facility and the McMaster Biosafety Office. M.S.M was supported, in part, by funding from ALS Canada, the Natural Sciences and Engineering Research Council of Canada (NSERC), the M.G. DeGroote Institute for Infectious Disease Research, and by a Canada Research Chair in Viral Pandemics, A.M. was supported by an Ontario Graduate Scholarship and an ALS Canada-Brain Canada Trainee Award.

## Author Contributions

Conceptualization, A.M., J.M., B.C., M.S.M.; methodology, A.M., J.M., B.C., M.S.M.; formal analysis, A.M., J.M., B.C., A.D.G.; M.S.M.; investigation, A.M., J.M., B.C., D.C., M.R.D, S.A., J.C.A., A.T.C, V.S., A.Z., H.D.S., M.L., Y.K., K.M., M.S.M.; writing, A.M., S.A., J.M., M.S.M.; supervision, M.S.M.

## Data and Materials Availability

All data, code, and materials reported in this article will be shared by the lead contact upon request.

