## Supplementary figures and images for "Acute Viral Infection Accelerates Neurodegeneration in a Mouse Model of ALS"

### Graphical Abstract

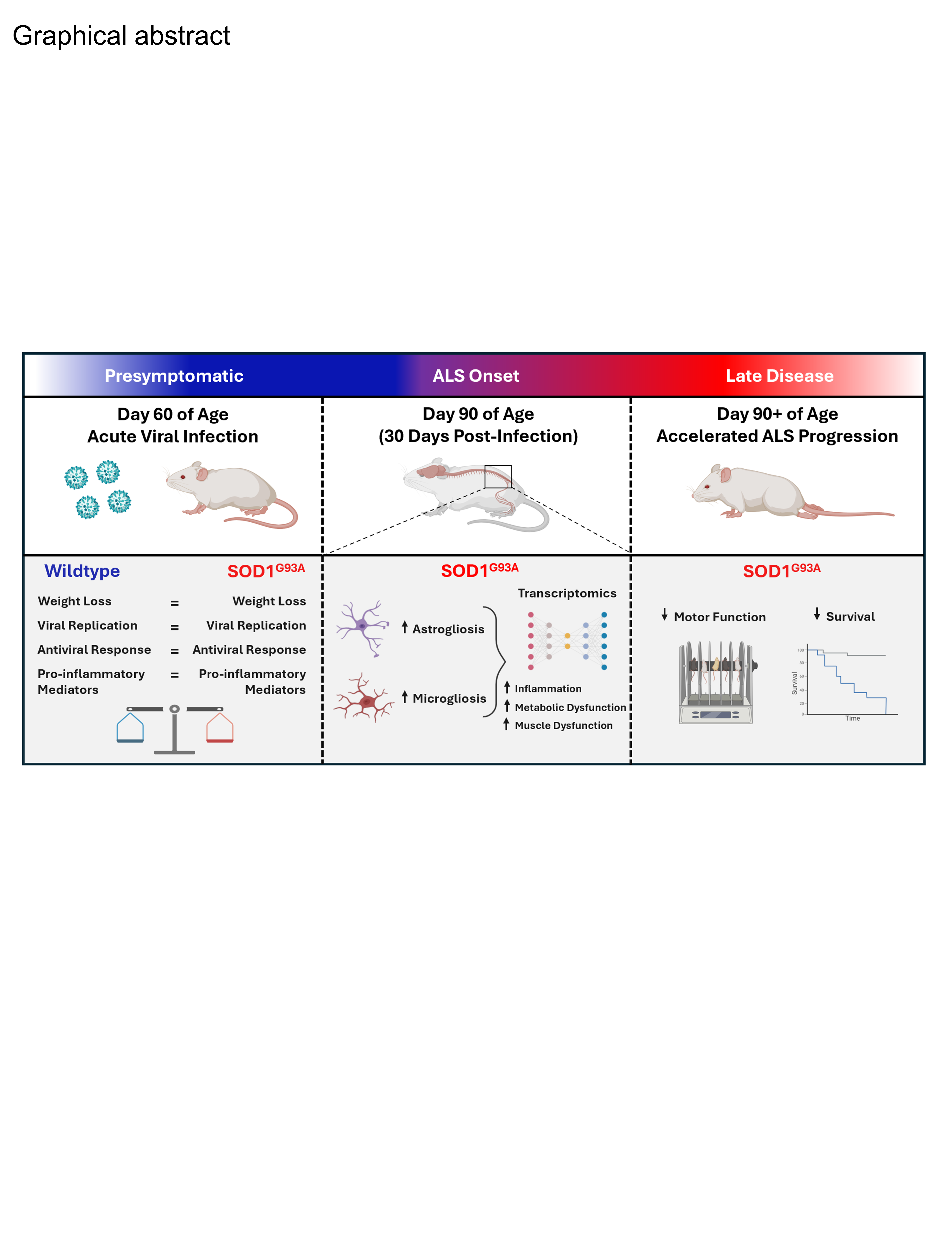

### Supplemental Figure 1

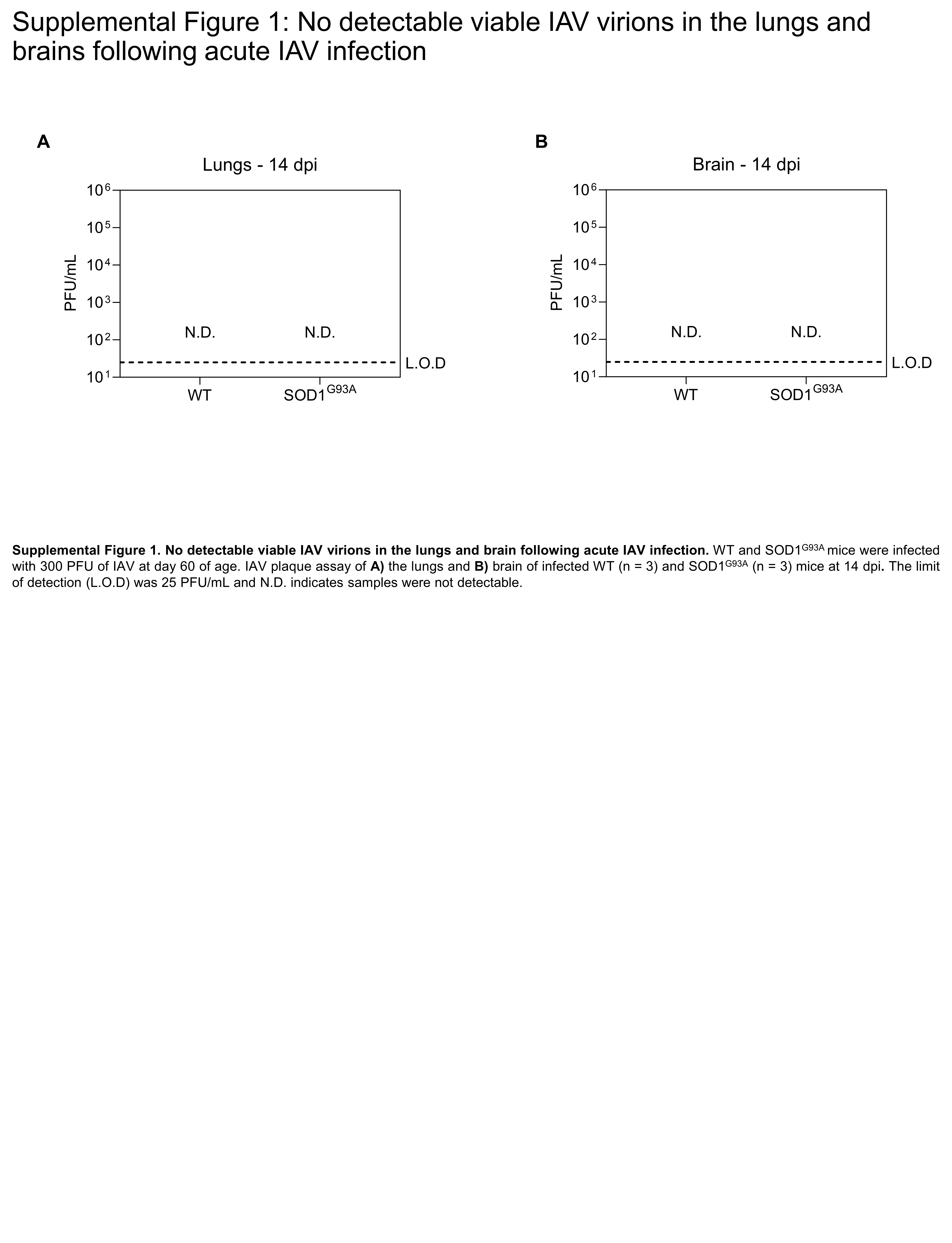

### Supplemental Figure 2

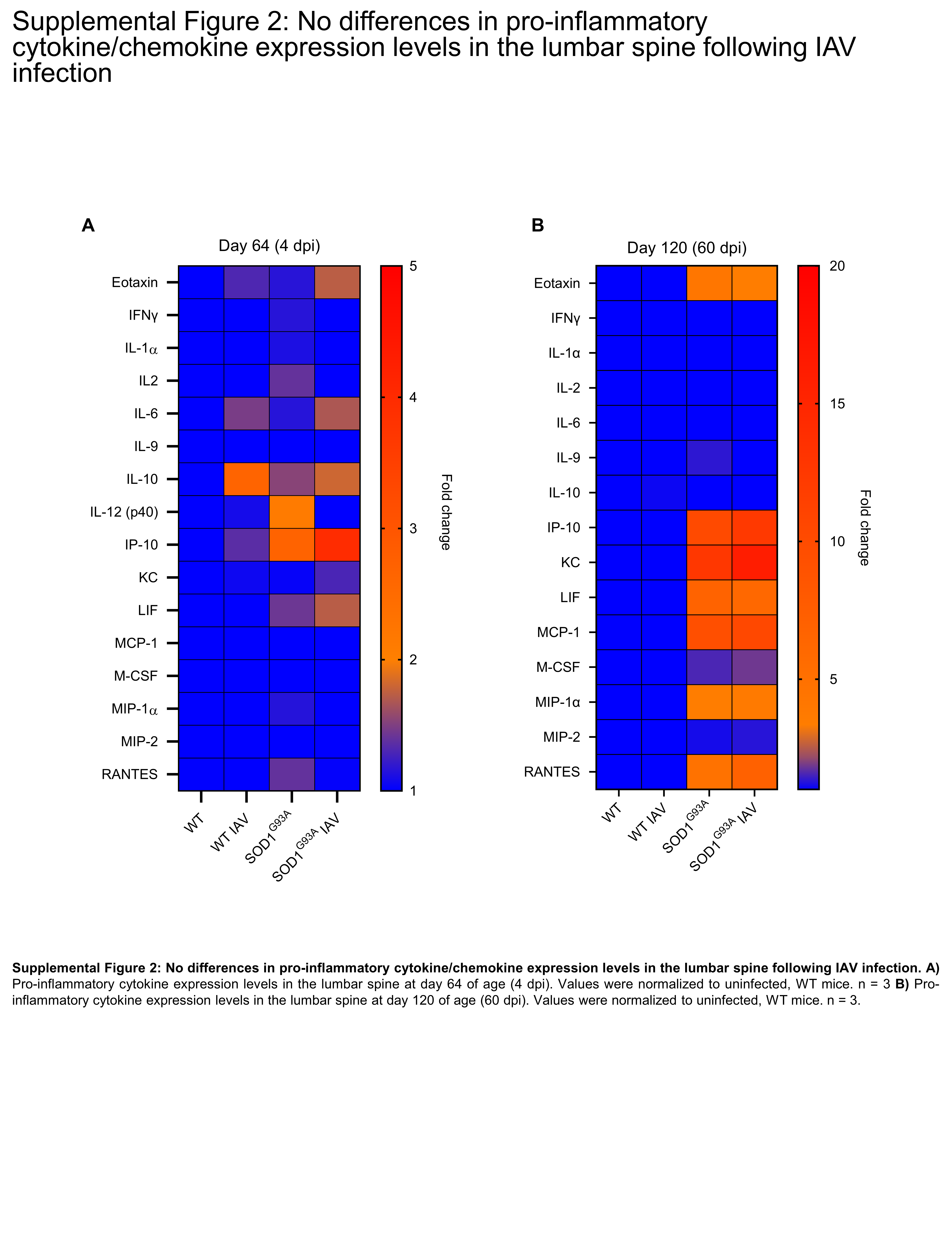

### Supplemental Figure 3

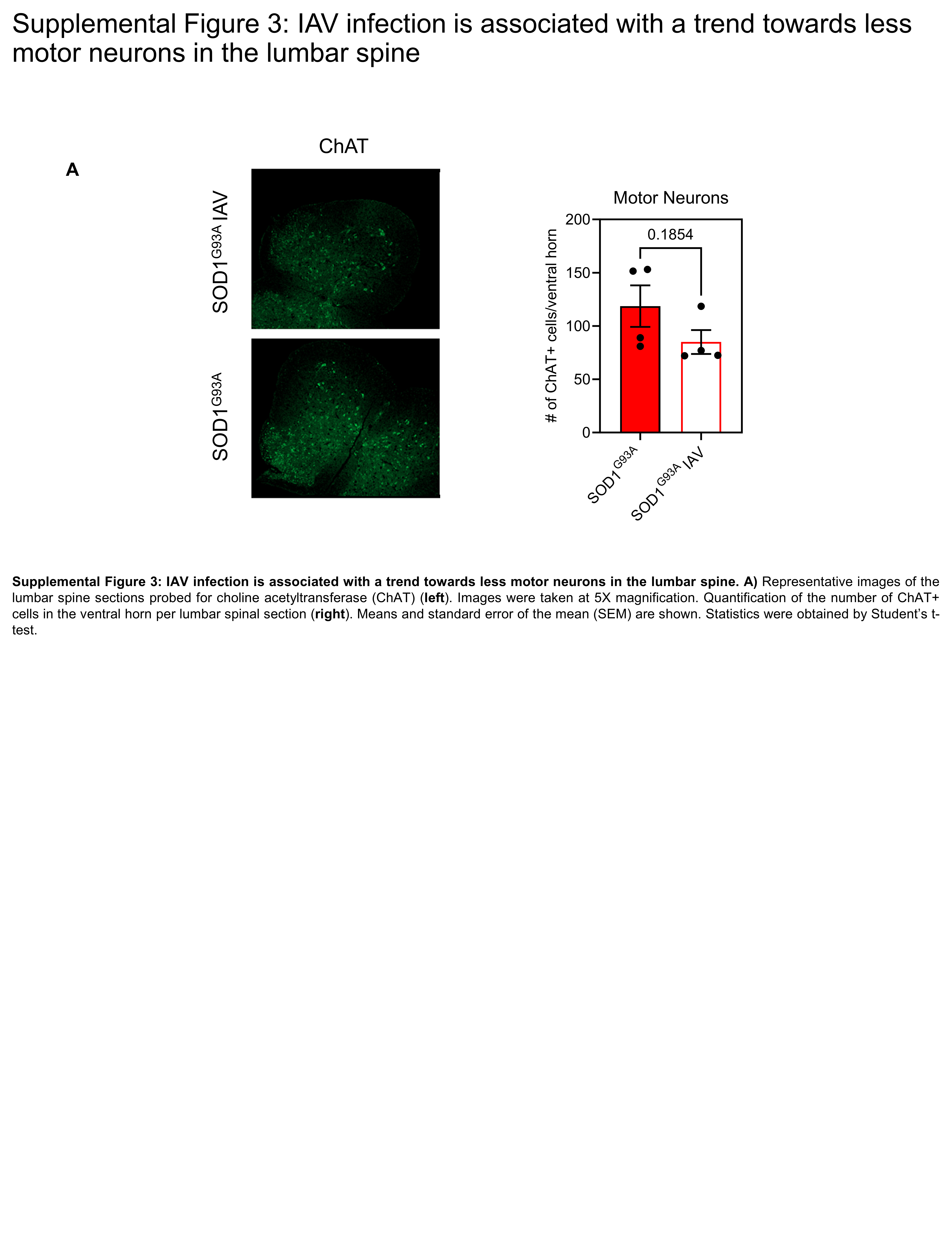
